# Single-AAV Delivery of the SpCas9 Nickase Improves CAG/CTG Repeat Contraction in a Mouse Model of Huntington’s Disease

**DOI:** 10.64898/2026.10.30.762787

**Authors:** Alvaro Murillo, Alys N. Aston, Mariah J. Lelos, Vincent Dion

**Affiliations:** UK Dementia Research Institute at Cardiff University, Hadyn Ellis Building, Maindy Road, Cardiff, United Kingdom, CF24 4HQ; School of Biosciences, Cardiff University, The Sir Martin Evans Building, Museum Avenue, Cardiff, CF10 3AX; Division of Psychological Medicine and Clinical Neurosciences, Cardiff University, Maindy Road, Cardiff, UK, CF24 4HQ

**Keywords:** CRISPR, AAV, Cas9 nickase, gene editing, Huntington’s disease, R6/1

## Abstract

The best studied Cas9 enzyme, from *Streptococcus pyogenes*, is thought to be too large to be efficiently packaged into a single adeno-associated virus (AAV) together with a single-guide (sg) RNA cassette, requiring a dual-vector system. Here we have circumvented this limitation and packaged the full CRISPR system into a single AAV by using a single promoter driving the expression of both the Cas9 enzyme and a sgRNA. We compared the ability of the dual and single vector systems to induce contraction of CAG/CTG repeats in the striatum of a Huntington’s disease model. We show that the all-in-one system led to larger contractions of the CAG/CTG repeats. This approach provides a more efficient and less costly approach for *in vivo* gene editing and improves the translational potential of AAV-mediated delivery of Cas9.

**One sentence summary:** Delivery of CRISPR-Cas9 nickase in a single AAV improves contraction of CAG repeats in HD.

## Introduction

Gene editing has been greatly facilitated by the advent of CRISPR. Specifically, the Type II CRISPR system from *Streptococcus pyogenes* has been the best studied so far. It contains a customizable CRISPR RNA (crRNA), which can be fused to a trans-activating RNA (tracrRNA) to produce a single-guide RNA (sgRNA), and the Cas9 (SpCas9) endonuclease^1,2^. This SpCas9 orthologue has been the subject of much improvement and modification, in large part because it has proven to be one of the most efficient Cas9 enzymes to date^3–6^. However, when it comes to *in vivo* gene editing, the SpCas9 has a significant drawback: it is composed of 1,368 amino acids (4.1kb worth of DNA)^2^, making it bulky for *in vivo* studies. Indeed, the most utilised gene delivery system, adeno-associated viruses (AAVs), can only hold up to about 4.7 kb of single-stranded DNA cargo^7^. The result is that two vectors must be used, one containing the SpCas9, the other the sgRNA^8,9^. This increases cost, reduces efficiency, and increases the risks for both *in vivo* and clinical applications^10^.

There have been multiple approaches tested to get around this problem. The first was to use a small Cas9 orthologue, most commonly from *S aureus (SaCas9)*, which is small enough to be packaged into a single vector together with its sgRNA^11^. SaCas9, however, is not as efficient at editing and the sgRNA target must be changed due to the difference in PAM sequence requirements. For some applications, therefore, this is not an option.

An entirely different approach is the use of lipid nanoparticles to encapsulate the SpCas9/sgRNA ribonucleoprotein complex^13^. Although promising, this approach lacks cell type specificity, sustained expression, and the ability to deliver these particles to the central nervous system remains limited^14^.

For approaches requiring high efficiency, specific PAM requirements, the targeting of specific cell types, and sustained expression, a different approach is necessary. One example of such an application is Cas9–nickase mediated contraction of the CAG/CTG repeats that cause expanded repeat disorders, including Huntington’s disease (HD)^9^. HD is a dominant hereditary neurodegenerative disorder characterized by cognitive and motor decline^15^. It is caused by the expansion of a CAG repeat in exon 1 of the Huntingtin (*HTT*) gene^16^ and disease-modifying therapies remain elusive^17^. The expansion results in transcriptome alterations, protein aggregation, and ultimately neuronal death, with the most susceptible cell type being medium spiny neurons^18^. Because the size of the repeat tract is the main modifier of disease outcomes^19^, contracting it is an attractive therapeutic option. We have previously demonstrated that we could contract the repeat tract efficiently by using the SpCas9D10A nickase with a sgRNA that binds the CAG/CTG repeat itself^20,21^. This works in both dividing and non-dividing cell types, at multiple disease loci, and it is free of detectable off-target mutations^9,21^. When injected, using two AAVs stereotactically into the striatum of the R6/1 HD mouse model, it partially rescued the transcriptome of medium spiny neurons, reduced the inclusion body burden, and improved some of the locomotor behaviours^9^. We have shown previously in this context that several Cas9 orthologues with different PAM requirements were unable to induce contractions^12^. Moreover, we require sustained expression of the Cas9 nickase as contractions continue to accumulate over time^9^. This is therefore a case that would benefit from being able to package the SpCas9 in a single AAV for *in vivo* delivery.

Here, we have created an all-in-one AAV construct that drives the *in vivo* expression of both the Cas9 nickase and the sgRNA from a single H1 promoter^22,23^. We show that intra-striatal delivery of the all-in-one construct is more efficient in contracting the repeat tract compared to the previous dual AAV approach in the R6/1 mouse model. Our results provide the first proof-of-concept for the use of a single AAV to deliver the full *S pyogenes* CRISPR-Cas9 system *in vivo*, overcoming an important hurdle towards the translation of *in vivo* gene editing approaches.

## Material and Methods

### Plasmids and sgRNA design

Two vector system plasmid constructs were described in Cinesi et al., 2016 and Murillo et al., 2026. All-in-one constructs were designed following the sequences described from Gao et al., 2019^23^ and adapted for our sgCTG_6 a_nd Cas9 nickase^9^.The tRNA sequence used in the dual CRISPR systems was previously described^24^ and incorporated into our plasmids. Guides RNA were designed using the ChopChop webtool^25^. The plasmids used are in Supplementary table S3 and sgRNA sequences in Supplementary table S4.

### Cell lines, culture conditions and transfection

GFP(CAG)_101 c_ells were maintained at 37°C with 5% CO_2 i_n Dulbecco’s modified Eagle’s medium (DMEM) glutamax, supplemented with 10% dialysed foetal bovine serum (FBS, Gibco), 100 U ml^−1^ penicillin, 100 μg ml^−1^ streptomycin, 15 μg ml^−1^ blasticidin and 150 μg ml^−1^ hygromycin. The media was refreshed every 2–3 days. GFP(CAG)_101 c_ells were previously genotyped using the Cell Line Authentication service from Eurofins and they were confirmed to be HEK293.2sus^21^. Mycoplasma testing was performed by Eurofins and was found to be negative.

Transient transfections of GFP(CAG)_101 c_ells were conducted as described^26^. Cells were cultured for up to 12 days in the presence or absence of 2 μg ml^−1^ of doxycycline (dox). Cells were collected 3 days after transfection for genomic DNA extraction for *POLQ*, *FAN1* and *GFP* amplification using the primers listed in Supplementary table 5, and 12 days after transfection for flow cytometry analysis.

### T7 endo I assay

For the T7 endo I assays, the targeted regions were amplified (see list of primers in Supplementary table 5). The PCR was done using the Mango Taq polymerase (Scientific Laboratory Supplies) and the cycles used were 95 °C for 5min, followed by 30 cycles at 95 °C for 30’’, 55 °C (*POLQ* and *GFP*) 53°C (*FAN1*) for 30’’ and 72 °C for 30’’ followed by a final step at 72 °C for 10 min. PCR products were purified using the DNA Clean and Concentrator kit (Zymo). The annealing of PCR products was performed by mixing 200 ng of DNA in 1 × NEBuffer. After annealing, 10 U of T7 endonuclease I (NEB) was added and incubated for 15 min at 37 °C. We determined genome targeting efficiency using the equation “% gene modification = 100 x (1 – (1-fraction cleaved)^1/2^)” previously published^27^. Band intensity quantification was performed with ImageJ (v2.3.051).

### Quantification of editing at the GFP transgene

GFP(CAG)_101 w_ere cultured in the presence of dox (2 μg ml^−1^) and transfected as above and quantified as previously^12,26,28^. Briefly, we first established gates that contained the top 1% of GFP-expressing cells transfected with the nickase plasmid only. For each experiment, the gates were adjusted accordingly. The percentage of cells from the test population falling in the GFP*^−^*population (Supplementary Fig. 1c) was then quantified. In preparation for flow cytometry analysis, cells were re-suspended in phosphate-buffered saline (PBS) with 1 mM EDTA to a concentration of about 10^6^ cells per ml. For each condition, we measured at least 2 × 10^5^ events using a Thermo Attune NxT Acoustic Focusing Cytometer. Data analysis was done using FlowJo software v.10.8.1. The sgRNAs (Supplementary Table 4) used for gene editing of the GFP transgene targeted exons.

### Quantification of contractions using the GFP-based reporter

Changes in CAG repeat length were approximated using a GFP-based reporter system as previously reported^21,28^. To quantify the fold increase in the number of GFP^+^ cells, we first draw gates that contain the top and bottom 1% of GFP-expressing cells in the control treatment, which in this case was cells transfected with the nickase plasmid only. For each treatment, the top and bottom 1% were adjusted to take any day-to-day shift in GFP expression into account. We then calculated the percentage of cells from the test population falling within these same gates.

### Long read sequencing of repeats

Expanded repeats were sequenced as before using SMRT HiFi amplicon sequencing^29^ (see Supplementary table 5 for primers). Mouse samples were mechanically disaggregated in Nucleospin Tissue DNA extraction T1 buffer using a mini hand-held homogenizer (ThermoFisher). Genomic DNA was extracted with the NucleoSpin Tissue DNA extraction kit (Macherey-Nagel). Three PCRs (3 x 50 µl) per sample were combined and cleaned with AMPure PB beads according to PacBio’s barcoded overhang adapter protocol (Pacific Biosciences #101-791-700). Barcoded sample pool purity was analysed on the 5200 Fragment Analyser (Agilent) and final library concentration determined using Invitrogen Qubit 1X HS dsDNA kit. Sample pools were sequenced using the Pacific Biosciences Sequel IIe.

Precise CAG/CTG repeat sizes were determined using Repeat Detector^29^ (version 1.0.15eb445). Unaligned reads were assessed using the restrictive profile with a repeat size range of [0–250]. For each analysis, the – with-revcomp option was enabled and data was output to a histogram (-o histogram option). The histograms were plotted using GraphPad Prism software (v.10.0.0). As the sequences are unaligned when we determine repeat size, the CCSs containing between 0 and 5 CAG/CTGs are removed from analysis as they include a large proportion of truncated PCR products and other sequences that do not align with the *HTT* locus.

### Area under the curve, repeat size indices, and comparisons

Area under the curve analysis was calculated as described^9^. In short, total reads were normalised and adjusted to the mode as 0. Then, we subtracted the frequency of reads of the treated samples from those in the control conditions for each repeat size to create delta plots and test the differences^30^. It represents the frequency of alleles that has changed compared to the control samples. The instability index was calculated as previously described in^9,31^. The expansion and contraction indices were calculated such that expansion index + contraction index = instability index.

### Mouse housing

C57BL/6J R6/1 mice^32^ (B6.Cg-Tg(HDexon1)61Gpb/J) were maintained as hemizygotes with *ad libitum* access to food and water and maintained in a temperature and humidity-controlled environment on a 12 h dark/light cycle. The R6/1 mice used here were of both sexes and had between 118 and 130 CAG/CTG repeats. Health and wellbeing of the animals were monitored daily, and their weight were checked every week in a T1 biosafety level 2 animal facility. All experimental procedures done in Cardiff followed protocols in accordance with the United Kingdom Animals (Scientific Procedures) Act of 1986. All experimental procedures performed on R6/1 mice were approved by Cardiff University Animal Welfare and Ethical Review Board and carried out under the Home Office License PP7595333.

### Stereotactic injections

Mice were anaesthetised with isoflurane and placed in a stereotactic frame. In adult (2-months old) mice, the scalp was shaved, a longitudinal incision was made to expose the skull surface, and 2 burr holes were drilled above the infusion sites. A 4 μl viral suspension (7.98x10^10^ vg all-in-one AAV) was stereotactically injected into the striatum of adult mice according to the Paxinos and Franklin mouse brain atlas (AP +0.8; ML +-1.8; DV -2.2 mm from bregma). Hamilton syringes 701 N 10 µl were used for the infusion in adult mice. The infusion rate was 200 nL min^-1^, and the needle remained in place for 5 min after infusion for vector absorption before removal. The injection site was closed with stitches, and mice were allowed to recover in incubators. The two-vector system conditions are described elsewhere^9^.

### Western blotting

To assess Cas9 levels in GFP(CAG)_101 c_ell cultures, proteins were extracted using RIPA buffer supplemented with cOmplete^TM^ EDTA-free protease inhibitors. Samples were incubated for 30 min in a rotator at 4°C and soluble fractions were collected by centrifugation (30 min; 13000g at 4°C). Protein concentrations were determined using Pierce BCA protein assay (ThermoFisher). 20 μg protein was loaded on 4-12% Bis-Tris Plus precast gels (ThermoFisher) and run in 1X MES buffer (ThermoFisher) alongside Bio-Rad Kaleidoscope molecular weight marker. Proteins were transferred onto a nitrocellulose membrane using the Bolt system (ThermoFisher). Membranes were incubated overnight with the anti-Cas9 (1:1000; Abcam; ab191468) and anti-actin (1:2000; Sigma; A5441) primary antibodies, followed by incubation with alexa-680-conjugated secondary antibodies (1:1000). The results were imaged on the Odyssey Imager (LI-COR Biosciences).

### Statistics

Changes in GFP^+^ cells were compared using a one-way ANOVA followed by a post-hoc Tukey’s multiple comparison test. A nonparametric paired Wilcoxon or a nonparametric Kolmogorov-Smirnov for unpaired samples were used for allele frequency distribution in histogram plots and delta plots in the *in vivo* work. Furthermore, differences in instability indices over time were calculated using a one-way ANOVA with a post-hoc Tukey’s multiple comparison test. We used two-way ANOVA to estimate differences on treatment and time, followed by a post-hoc Tukey’s multiple comparison test. For repeat size comparison, in allele frequency or delta plot analysis, after long-read sequencing two tests were used. For comparison of repeat instability between striatal samples within the same animal a paired nonparametric Wilcoxon signed rank test was used. Statistical analyses were done using GraphPad Prism software version 10.0.0 and were always two-tailed tests using a significance cut off at P=0.05.

### Q-PCR

Quantitative real-time PCR (qPCR) was performed using the Applied Biosystems QuantStudio 7 Flex Real-Time PCR system and analysed with the QuantStudio Real-Time PCR Software (v1.7 Thermo Fisher). Briefly, qPCR was assayed in a total volume of 10 μl reaction mixture containing the ready-to-use FastStart Universal SYBR Green Master (ROX) (Thermo Fisher) and 0.5 µM of forward and reverse mix appropriate primers (Supplementary Table 5). All qPCR reactions were run in triplicates. Mean cycle threshold (Ct) values for each reaction were recorded, and the relative DNA copies were calculated and normalised to Actin (2^(ActinCt-geneCt)).

### Data availability statement

The CCSs sequences are available from SRA (https://www.ncbi.nlm.nih.gov/sra) PRJNA1483968.

## Results

### Efficient gene editing of an all-in-one construct in human cell lines

We first investigated the possibility of using an all-in-one construct for gene editing in immortalised cell lines. To address this, we generated a construct using a design first proposed by Gao et al^23^. It relies on the use of a single H1 promoter that leads to a Pol III transcript containing the sgRNA and a Pol II transcript for SpCas9 expression^23^. We have produced the same construct, but expressing the SpCas9D10A nickase (Fig. 1a). We used two sgRNAs that target opposite strands in close proximity, creating a double-stranded break for error-prone repair and the production of insertion/deletion mutations. In this construct, the sgRNAs flank a tRNA that is spliced out upon transcription^24^. The construct remains small enough to be packaged into a single AAV (Fig. 1a). We transfected both constructs in HEK293-derived GFP(CAG)_101 c_ells to edit *POLQ*, *FAN1* and the ectopic *GFP* present on chromosome 12 in this line^33,34^ (Fig. 1b). After transfecting a plasmid containing the single H1 promoter, the sgRNAs and the Cas9 nickase, we found efficient editing efficiency at all three loci using a T7 endonuclease I assay (Fig. 1cd). Importantly, the all-in-one CRISPR Cas9 nickase system has a similar genome editing efficiency as the classical Cas9 nuclease, despite requiring two simultaneous DNA nicks on opposite strands from two different sgRNAs. The constructs led to 39% and 36% targeting of *POLQ* as well as 18% and 23% editing of *FAN1* for the nuclease and nickase constructs, respectively (Supplementary Fig. 1a, Fig. 1e). However, the nickase construct showed a 2-fold decrease in editing efficiency at the GFP locus compared to the Cas9 nuclease (Fig. 1d), perhaps because the additional sgRNA used with the Cas9 nickase is inefficient. We confirmed these results using GFP intensity as an approximation of the number of cells mutated by each approach (Supplementary Fig. 1bc, Fig. 1e). These results suggest that both approaches within the all-in-one constructs can be used for knockout generation.

**Figure 1.**
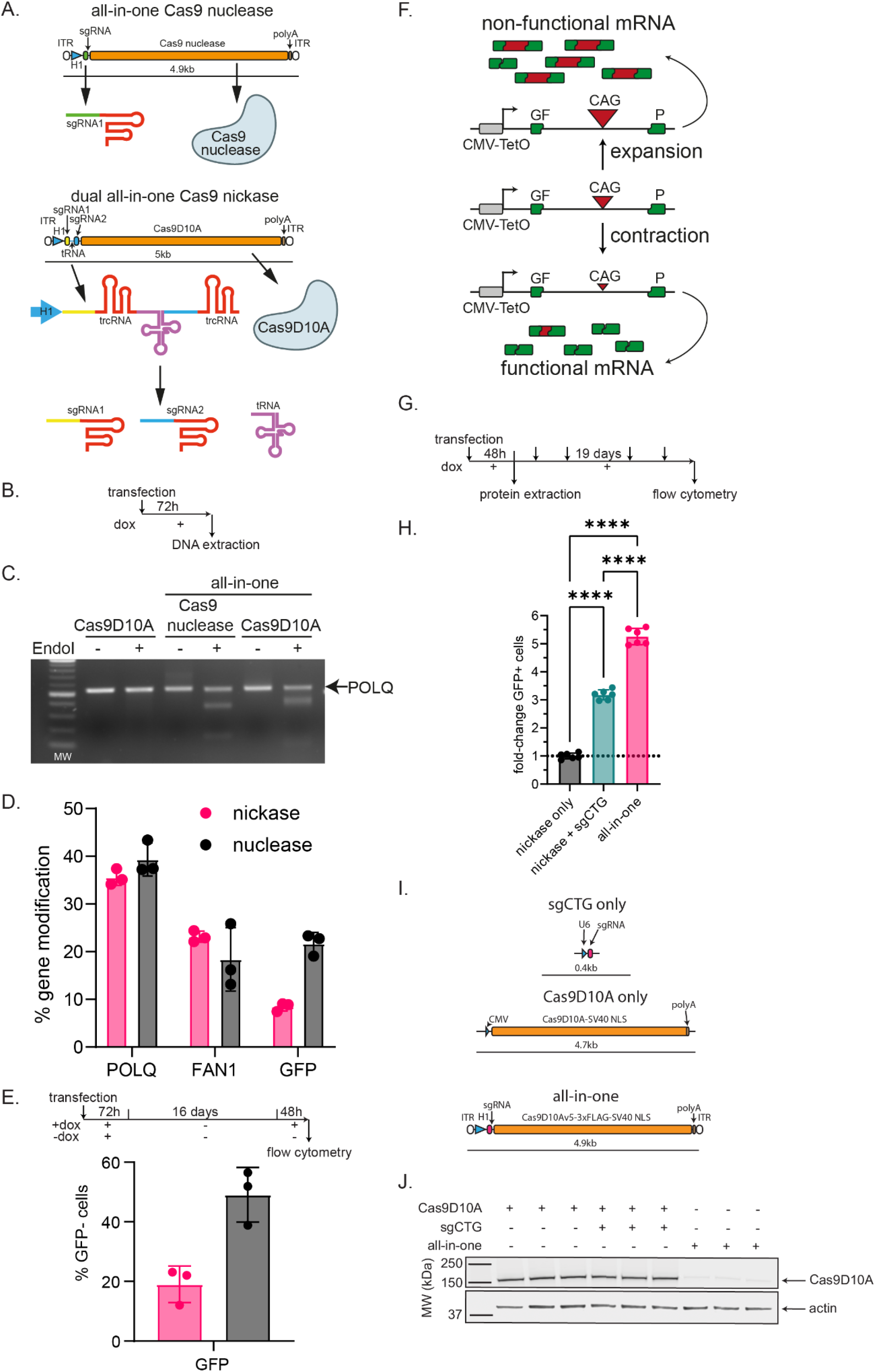
All-in-one constructs induce mutagenesis in several loci. A) Schematic representation of the all-in-one constructs. All plasmids are in a pAAV backbone and include the ITRs. All-in-one constructs expressing the Cas9 nuclease or the Cas9 nickase and a sgRNA under a single H1 promoter. The nickase construct expresses two sgRNA separated by a tRNA that can be spliced out^24^. B) Timeline of the experiment C,D. C) Example of a T7 endo I assay targeting POLQ in GFP(CAG)101 cells. Full blots are found in Supplementary Fig. 4. D) Quantification of the T7 endo I assay. All the images are found in Supplementary Fig. 1a. E) Experimental timeline and quantification of the percentage of GFP^-^cells as measured by flow cytometry (Supplementary Fig. 1bc). Note that the doxycycline (dox) is there to turn on transcription through the repeat tract, allowing Cas9 to access the DNA^21,26^. F) GFP-based assay used to detect contraction of the CAG repeats (modified from^20^). G) Experimental timeline of the experiment in I and J. H) Comparison of GFP^+^ cells after transfection of each construct. Dashed lines represent the GFP+ cells in the population with Cas9D10A only. One way ANOVA, ****P<0.001, post-hoc Tukey’s multiple comparison test. Error bars represent ± standard deviation. I) All-in-one construct for targeting the CAG/CTG sequence. J) Western blot of Cas9D10A and actin expression from GFP(CAG)101 cultures transduced with pcDNA3.3-TOPO_Cas9D10A nickase, pPN10_sgCTG, or pAAV_H1_sgCTG-Mut_ Cas9D10A_Muscle codon Optimized 48 hours after transfection. Full blots are found in Supplementary Fig. 4.

### All-in-one vector induces CAG repeat contractions in a GFP-based assay

We then assessed whether the all-in-one construct could induce contractions of the ectopic CAG/CTG repeats in GFP(CAG)_101._ The ectopic repeat in this line is found within the intron of a GFP mini gene. This reporter works because CAG repeats interfere with the splicing of the GFP mini gene in a size-dependent manner, with longer repeats producing less GFP^28,34^ (Fig. 1f). Therefore, we can use GFP intensity as a proxy for repeat size^20,21,28,34^. We transfected the all-in-one construct and measured GFP levels 12 days after the first transfection by flow cytometry (Fig. 1gh). We found a 1.65-fold increase in GFP^+^ cells when comparing the all-in-one vector to a two-vector approach, suggesting that having both the Cas9 nickase and sgCTG on the same plasmid increases efficiency (Fig. 1i). Importantly, this increase in efficiency was seen despite a lower expression of the Cas9 nickase (Fig. 1j). We conclude that a single promoter driving the Cas9 nickase and the sgCTG robustly induces contractions of CAG/CTG repeats in this cell line.

### Efficient editing of expanded CAG/CTG repeats in the R6/1 mouse striatum upon delivery of the all-in-one AAV

Next, we aimed to determine whether the all-in-one could edit DNA *in vivo*. To this end, we packaged our 4.6kb cargo into a single AAV serotype 9, totalling 4.9kb with the ITRs. We have shown previously that this serotype preferentially transduces medium spiny neurons^9,35^. Adult R6/1 mice (2-months old) were stereotactically injected with the all-in-one AAV into one striatum and the contralateral striatum received the injection of an AAV-sgCTG-GFP, as a control (Fig. 2ab). As in the previous study, we found robust infection efficiencies of the all-in-one AAV with an average of 734.9 ± 206, 843.9 ± 213, and 1011 ± 618 AAV DNA copies 1-, 2- and 3-months post-injection respectively (Supplementary Fig. 2). For comparison, our previous injections using a two-vector system contained 330 ± 92 for the Cas9 nickase AAV and 1020 ± 173 for the sgCTG AAV at 3 months post injection, for a total of 1350 ± 196 AAVs copies (Supplementary Fig. 2). Therefore, we injected a slightly lower amount of AAVs using the all-in-one vector.

We assessed CAG repeat lengths 1-, 2- and 3-months post-injection by long-read PacBio sequencing as before (Fig. 2b-e). We confirmed contraction of the repeat in the *HTT* transgene, increasing in frequency and size over 3 months post-injection (Supplementary Table 2; Supplementary Fig. 3). This is also accompanied by an apparent reduction in expansions, which may be contractions or the prevention of somatic expansions (Fig. 2c). At 2- and 3-months post-injection there is a progressive reduction in CAG repeat lengths (Fig. 2de), showing larger differences in repeat size distribution between hemispheres at the latest timepoint (Fig. 2f).

**Figure 2.**
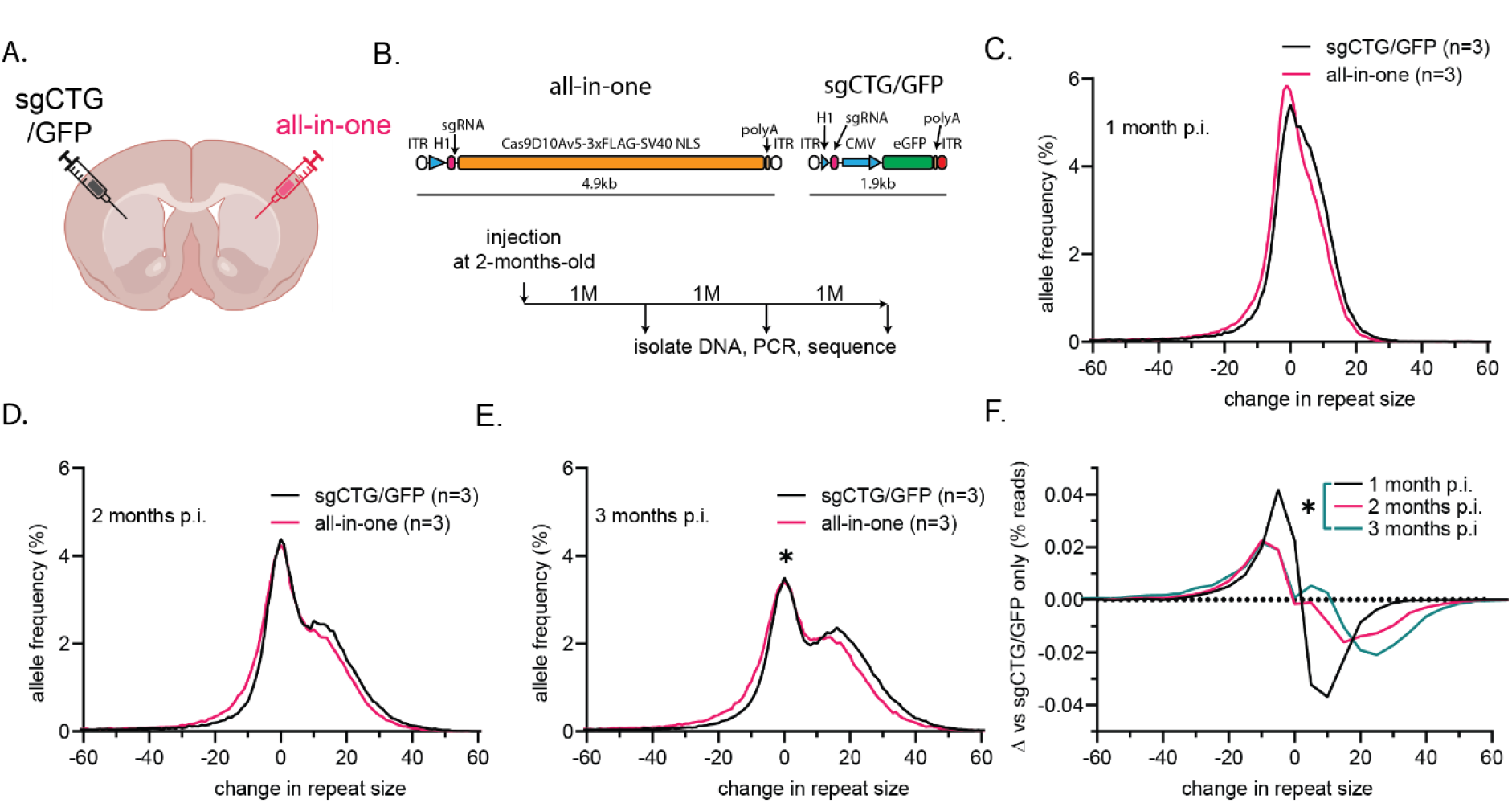
All-in-one AAV induces CAG repeat contraction in adult-injected R6/1 animals. A) Schematic of the experimental approach. B) Top: Schematic of the AAV constructs used. Bottom: timeline of the experiment. C) Aggregated graphs of repeat size distribution in hemispheres of R6/1 mice injected with sgCTG/GFP AAV only (black) versus all-in-one AAV (magenta) 1 month, D) 2 months and E) 3 months post-injection. *: P< 0.05 using a nonparametric Wilcoxon matched-pairs signed rank comparing repeat size in the all-in-one injected versus sgCTG-GFP injected only. F) Aggregated delta plot of repeat size distribution in hemispheres of R6/1 mice injected with all-in-one AAV subtracting sgCTG/GFP AAV repeat size distribution at 1, 2, and 3 months combined time points to show changes over time. Unpaired nonparametric Kolmogorov-Smirnov test was used to compare cumulative distributions (**P<0.01). Number of animals: n=3 per timepoint.

To benchmark the all-in-one vector, we compared it to our previous dual AAV system^9^. In the dual AAV constructs, we had the Cas9 nickase under a miniCMV promoter in one AAV9 and a second AAV9 expressing the sgCTG under a U6 promoter. The control hemispheres were injected with only the Cas9 nickase AAV. We found that the all-in-one AAV induced larger contractions of the CAG repeats and fewer expansions at 2- and 3-months post-injection compared to the 2-vector system (Fig. 3b-d). We also generated delta plots^30^ to compare the differences in repeat size between two treatments, which allow us to normalize each approach with its own control hemisphere. This analysis revealed significant differences in the contractions induced between treatments at 2- and 3-months post-injection (Fig. 3e-g). Those changes were mainly to the size of the contractions rather than the overall efficiency with the all-in-one construct inducing contractions of 19.7 repeats over 3 months (Supplementary Table 2). Analysing the data using instability indices instead showed a similar time-dependent trend of fewer expansions and more frequent contractions (Supplementary Fig. 3b-d).

**Figure 3.**
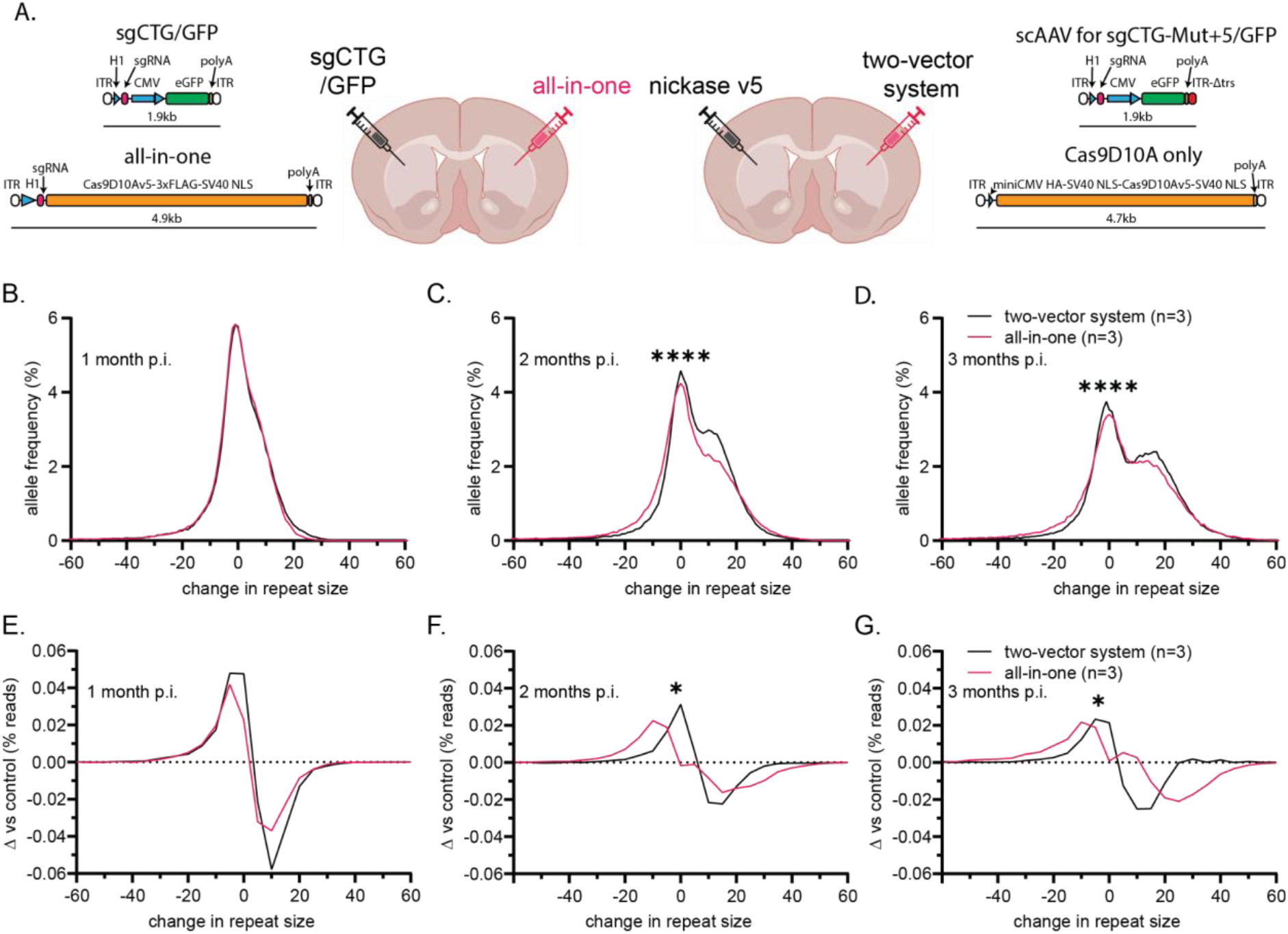
All-in-one AAV injection induces larger CAG repeat contraction in adult injected R6/1 animals compared to a dual vector system. A) Schematic of the experimental approach and of the AAV constructs compared. B) Aggregated graphs of repeat size distribution in hemispheres of R6/1 mice injected with the two-vector system (black -from ^9^) versus all-in-one AAV (magenta) 1 month, C) 2 months and D) 3 months post-injection. Note that the graphs are normalised to the modal peak size found in each hemisphere (see methods). Unpaired nonparametric Kolmogorov-Smirnov test was used to compare cumulative distributions (****: P<0.0001). E) Aggregated delta plot of repeat size distribution comparing the all-in-one AAV (magenta) to the dual-AAV injection (black) at 1 month, F) 2 months, and G) 3 months.Unpaired nonparametric Kolmogorov-Smirnov test was used to compare cumulative distributions (*: P<0.05).

Additionally, the instability index showed a slower increase over time in the hemisphere injected with the all-in-one vector compared to having two AAVs (Supplementary Fig. 3b; Supplementary Table 1), which was accompanied by a more negative contraction index (Supplementary Fig. 3c; Supplementary Table 1), and a lower expansion index (Supplementary Fig. 3d; Supplementary Table 1). Together, these results show that the all-in-one single vector system led to larger contractions *in vivo* compared to the dual AAV approach.

## Discussion

Here we showed that a single H1 promoter driving the expression of both a Cas9 and a sgRNA leads to efficient editing *in vitro*, including the ability of the Cas9 nickase to induce contractions of CAG/CTG repeats. This construct design allowed us to package the large SpCas9 and a sgRNA into a single AAV and obtain robust editing. This was done in the context of a HD mouse model, but is likely to be widely applicable, beyond editing CAG/CTG repeats.

Interestingly, the all-in-one AAV induced larger contractions, rather than a higher efficiency of gene editing. This was unexpected because once a cell has been infected with both the Cas9 nickase and the sgCTG, whether because they are dually infected or they are infected by the all-in-one vector, the enzymes are the same and should induce contractions in a similar manner. The observation is also not easily explained by the infection efficiency as the cells had an excess of viral genomes in both cases and thus the majority of the medium spiny neurons in the striatum are infected. It is perhaps because of the ratio of Cas9 to sgCTG produced from the H1 promoter. Indeed, we previously found that decreasing the Cas9 to sgCTG ratio to 1:2 improved contraction efficiency, but we could not go lower due to the AAV concentrations available^9^. However, Gao et al found that in human cells the H1 promoter expressed 1 Cas9 mRNA for every 3 to 4 sgRNAs^23^. Thus, we hypothesise that an all-in-one infected cell goes through multiple cycles of contractions once it is infected as a result of a better Cas9 to sgRNA ratio.

There are two more advantages to this all-in-one construct. First, the Cas9 nickase has a lower expression, despite a similar editing efficiency. Decreasing the amount of Cas9 has been shown to reduce off-target mutations^36,37^, which could lead to safer *in vivo* gene editing^38,39^. Moreover, we needed fewer viral genomes for the all-in-one construct, which is expected to decrease the potential risks of an immune response and reduce production cost.

Our approach is not without limitations. For example, gene editing requiring Cas9 fusion proteins, including base editing^40,41^ and prime editing^42^, would still be too large for packaging and require a dual AAV system. Nevertheless, the single-AAV delivery of SpCas9 has some major advantages that will help the clinical translation of *in vivo* gene editing-based treatments.

## Acknowledgements

Brain slices (Fig. 2-3) were created with BioRender.com (agreement number JM28EFZ3G9). We would like to thank Andrew Jefferson for training, Joanne Morgan and the core team at the Centre for Neuropsychiatric Genetics and Genomics at Cardiff University for support with PacBio sequencing, and Ruban Rex Peter Durairaj for support with the Repeat Detector run. Also, we would like to thank members of the Dion lab for critical reading of the manuscript.

## Author contributions

AM: Conceptualization, Funding acquisition, Project administration, Formal Analysis, Investigation, Methodology, Visualization, Writing – original draft, Writing – review & editing.

ANA: Methodology, Writing – review & editing

ML: Supervision, Writing – review & editing

VD: Conceptualization, Funding acquisition, Project administration, Supervision, Visualization, Writing – original draft, Writing – review & editing

## Supplementary Material

**Supplementary Figure 1.**
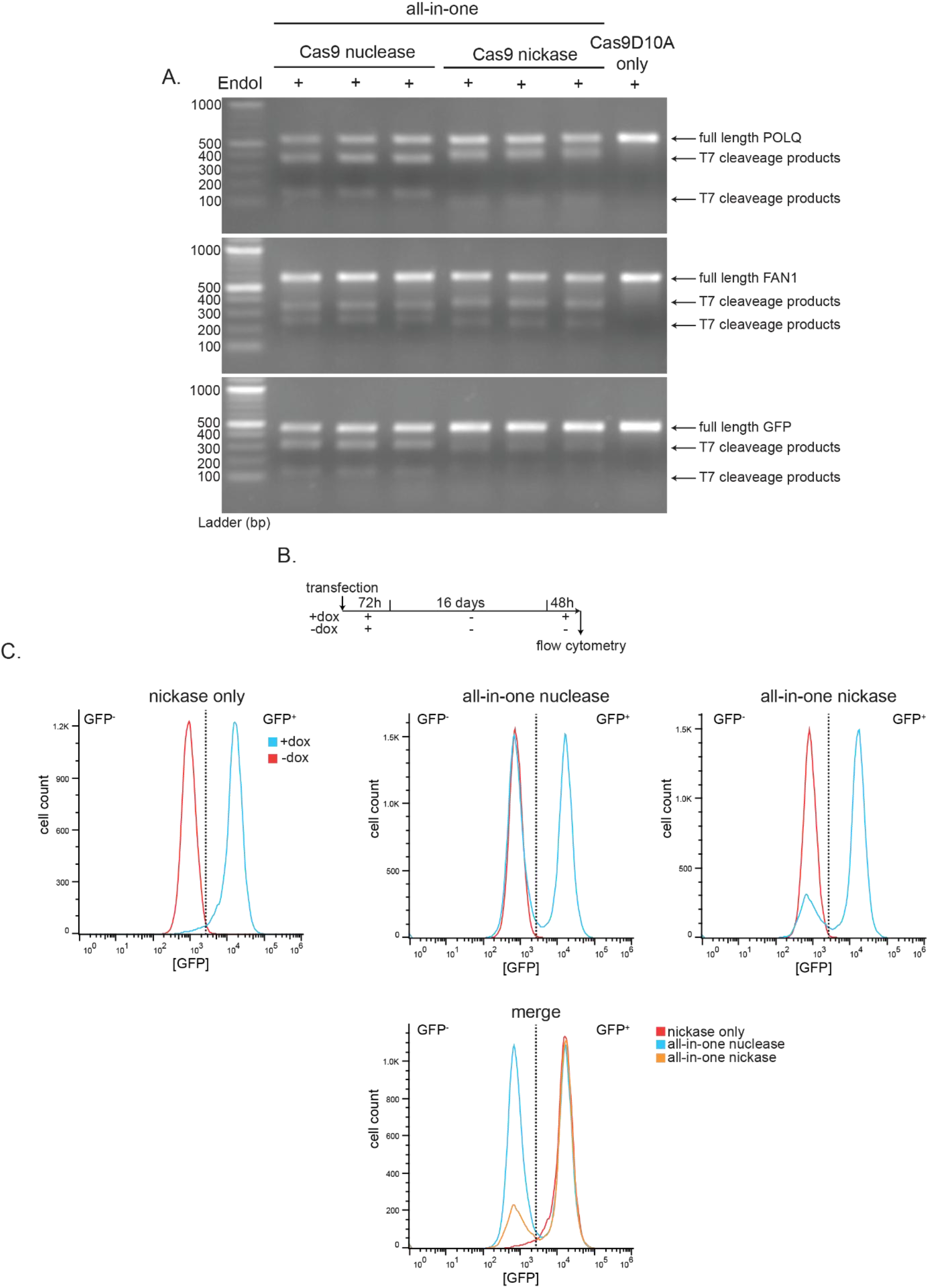
**The** all-in-one construct induces mutagenesis at several loci and contractions in CAG repeats. A) T7 endo I assay results targeting POLQ, FAN1 and GFP with the Cas9 nuclease and the CRISPR-Cas9D10A approaches Transfected GFP(CAG)101 with Cas9D10A only used as control. Three independent samples were collected and analysed per condition. Full blots are found in Supplementary Fig. 4. B) Timeline of the experiment shown in C. C) Representative flow cytometry profiles after expression of nickase only, all-in-one nuclease and all-in-one nickase approaches targeting GFP transgene in GFP(CAG)101 21 days after transfection. Dotted line represents the threshold for the top 1% of GFP-expressing cells in the nickase plasmid transfected only.

**Supplementary Figure 2.**
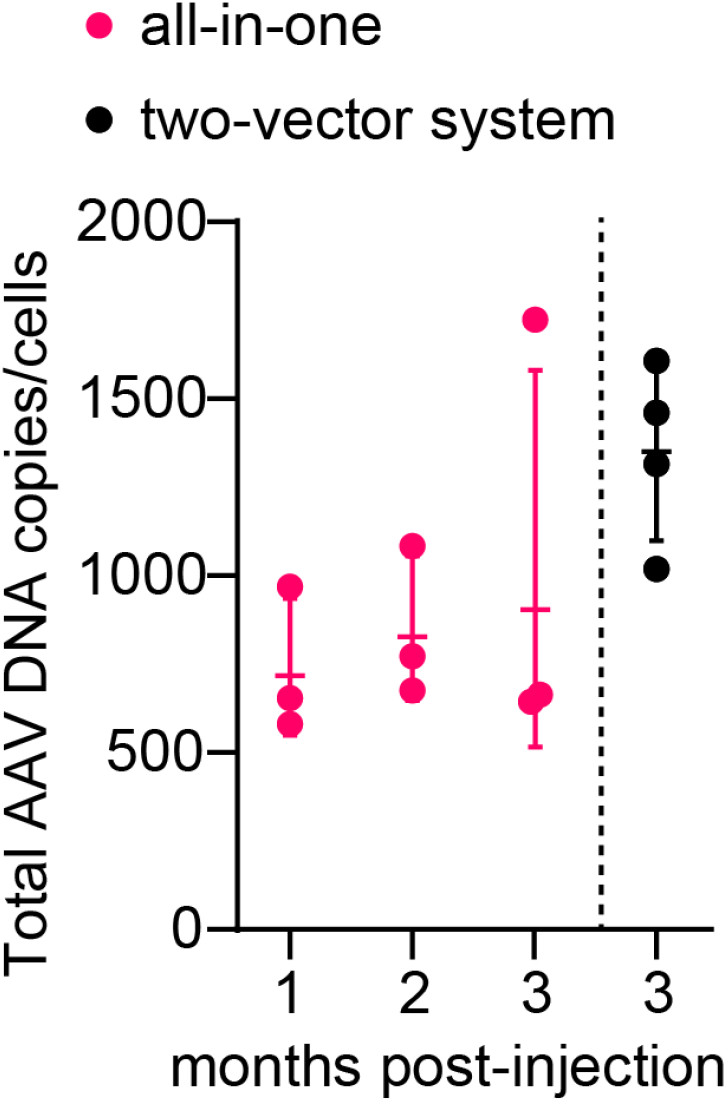
Number of AAV genome equivalent in the mouse brain. DNA extracted from striatal samples were used to determine the number of AAV genome copies by qPCR in treated mouse samples. AAV all-in-one copies were identified targeting nickase sequence. Data are mean ± SD (n = 3 animals per time point).. AAVs copies in mice injected with AAV-Cas9D10A v5 and scAAV-sgCTG-Mut+5 were obtained by the sum of both AAVs from the data published in Murillo et al., 2026^9^. Data are mean ± SD (n = 4 animals per condition).

**Supplementary Figure 3.**
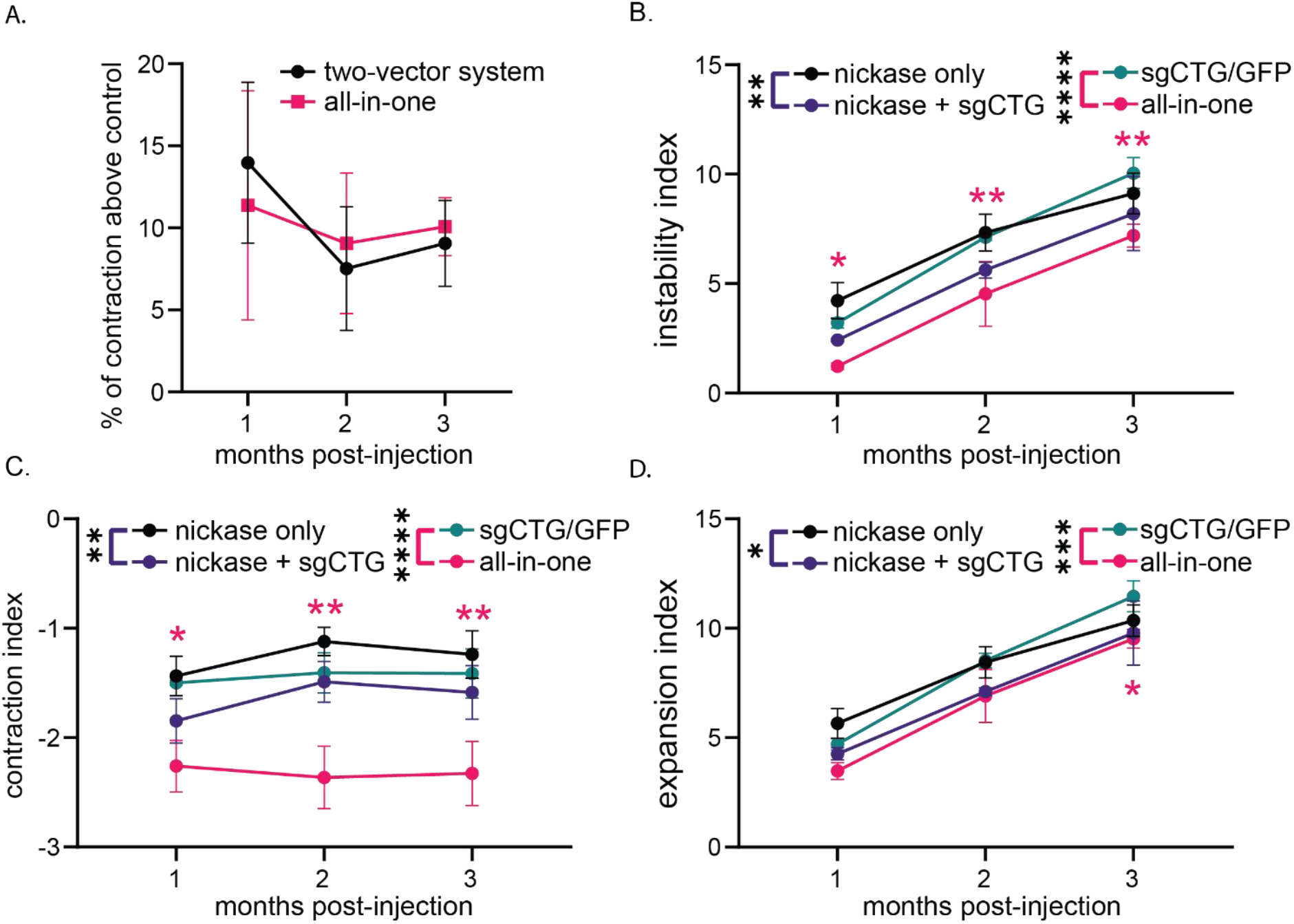
The all-in-one vector reduces instability and increases contraction index in comparison with a two vector system. Previously, the two vector system was published^9^. Difference in the area under the curve between the Cas9D10A v5 + sgCTG-Mut+5 ratio 1:2 (black) and all-in-one treatment (magenta). B) Instability index of the mice treated with the Cas9D10A v5 + sgCTG-Mut+5 ratio 1:2 (purple) versus Cas9 v5 only (black) and instability index of the mice treated with the all-in-one (magenta) versus sgCTG/GFP (green). C) Same as in B but showing only the expansion index and D) contraction index. A-D) Cohorts show an effect of time post-injection and the treatment using a two-way ANOVA (*P<0.05, **P<0.01, ***P<0.001, ****P<0.0001; magenta: all-in-one vs sgCTG/GFP; purple: Cas9D10A v5 + sgCTG-Mut+5 ratio 1:2 vs Cas9D10A v5 only). Two-way ANOVA, *P<0.05, **P<0.01, ***P<0.001, ****P<0.0001 post-hoc Tukey’s multiple comparison test. Error bars represent average ± standard deviation. The number of animals is indicated in Fig. 3.

**Supplementary Figure 4.**
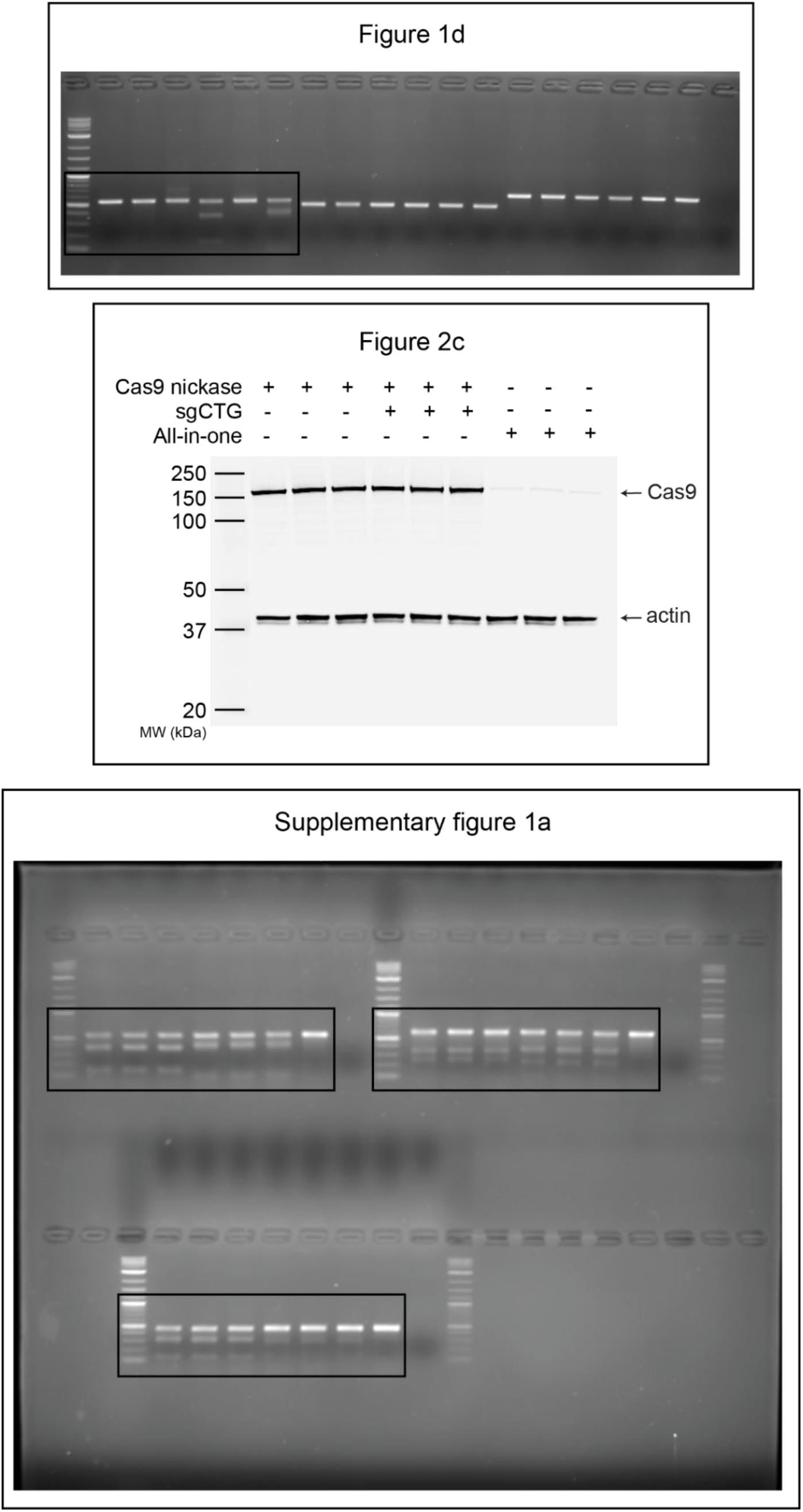
Unaltered full agarose gels and western blots membrane images. Black boxes indicate where the blots were cropped.

**Supplementary Table 1:**
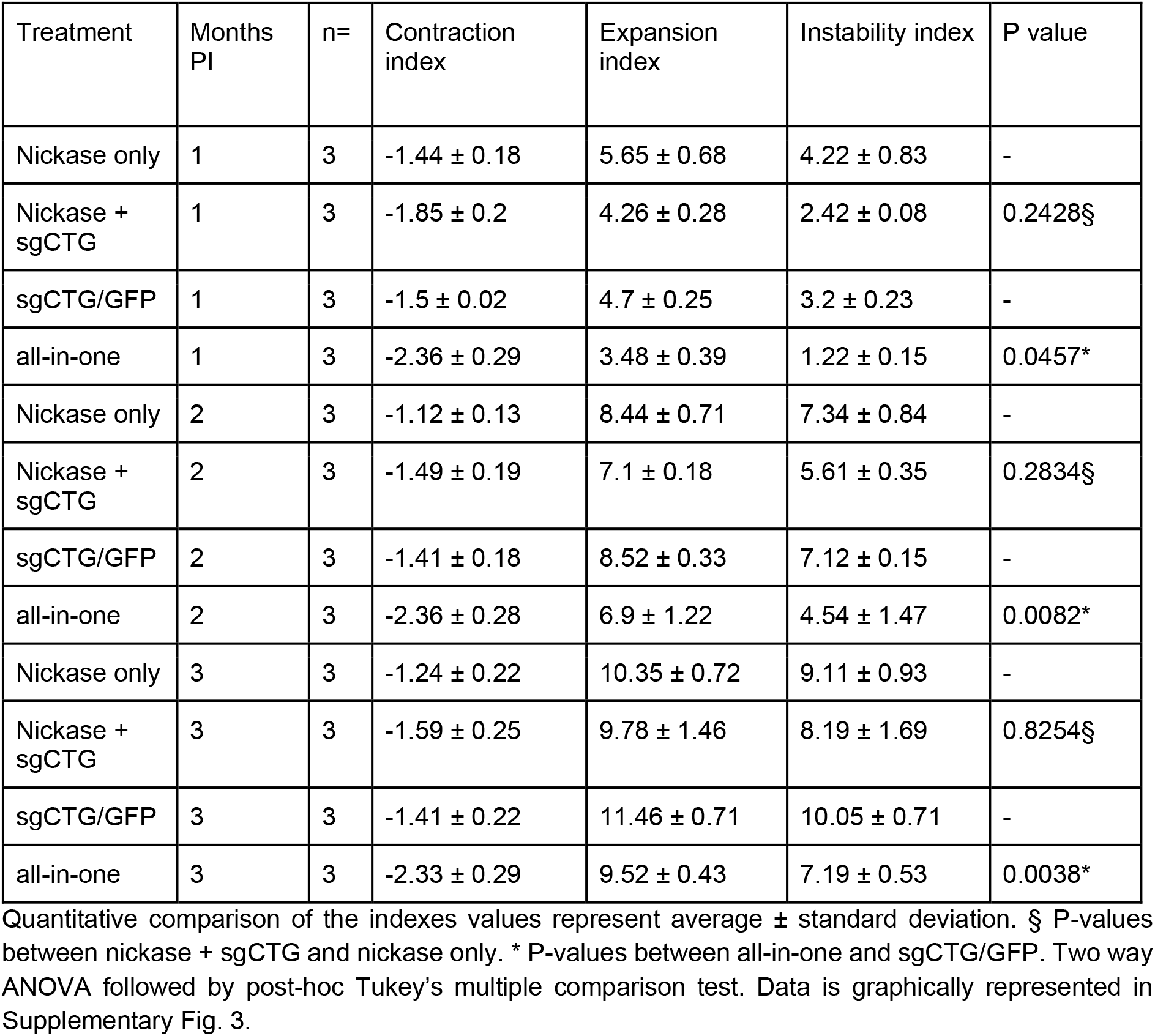
Instability indices in Supplementary Fig. 3.

**Supplementary Table 2:**
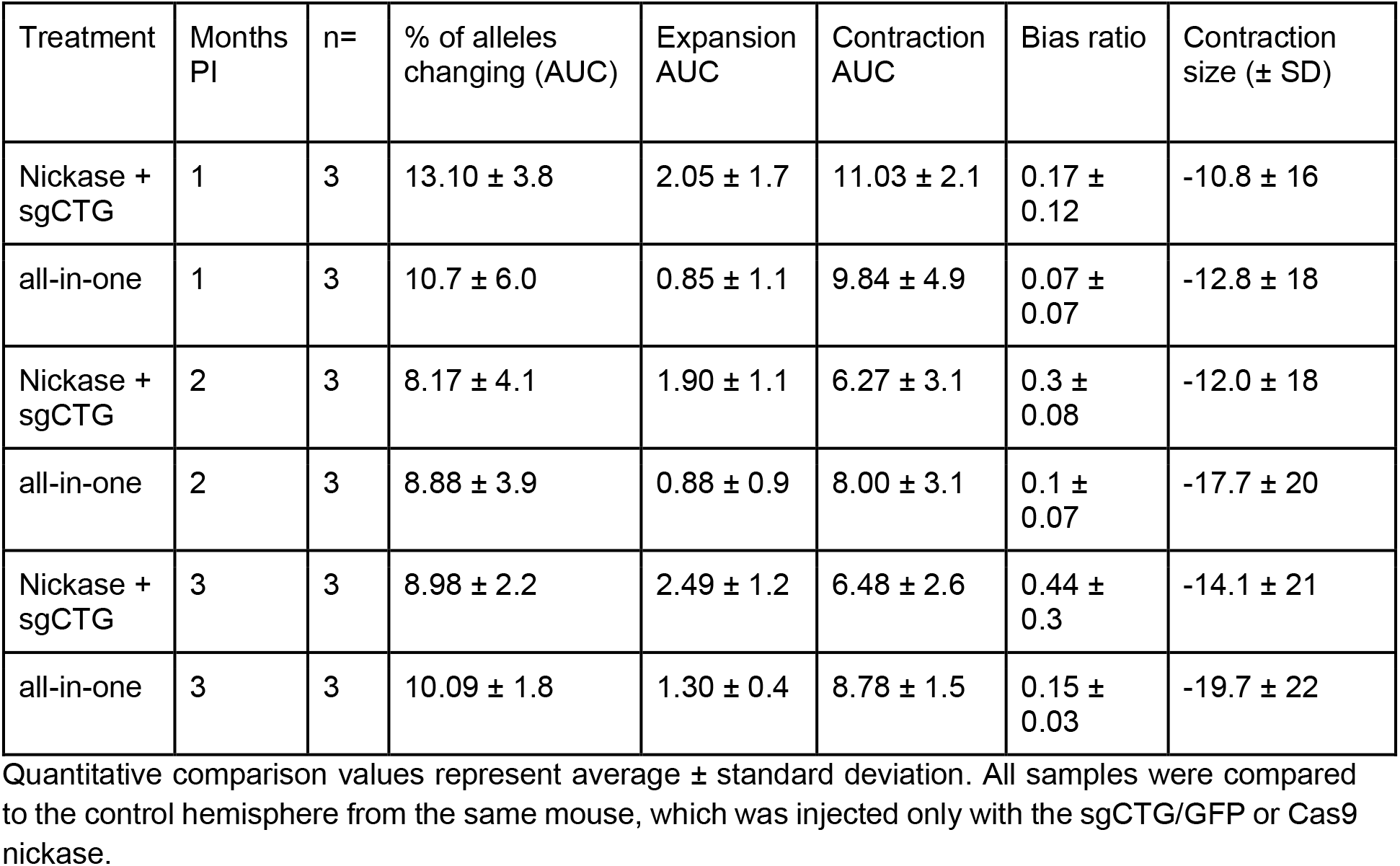
Instability statistics in Fig. 3.

**Supplementary Table 3:**
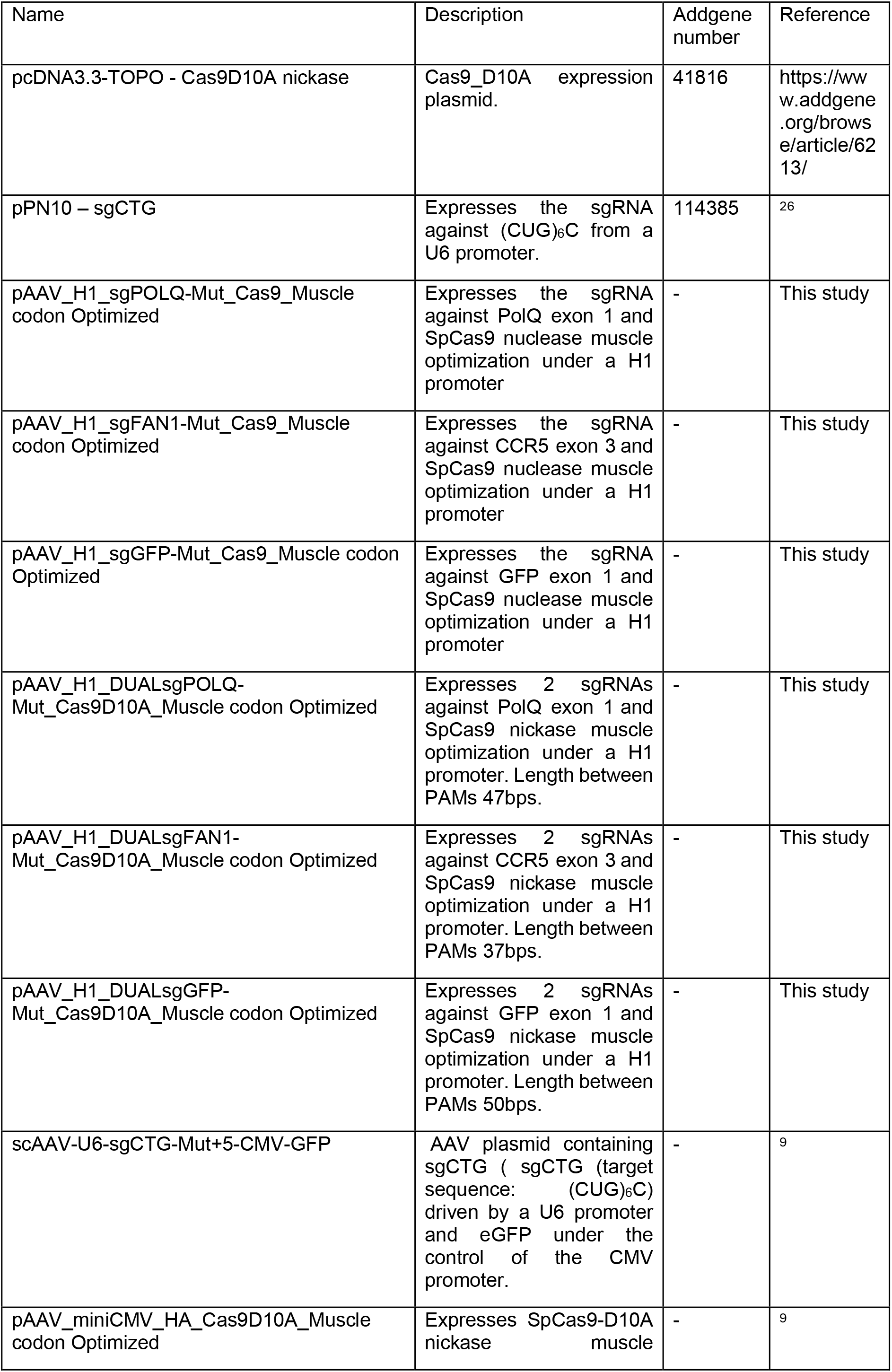
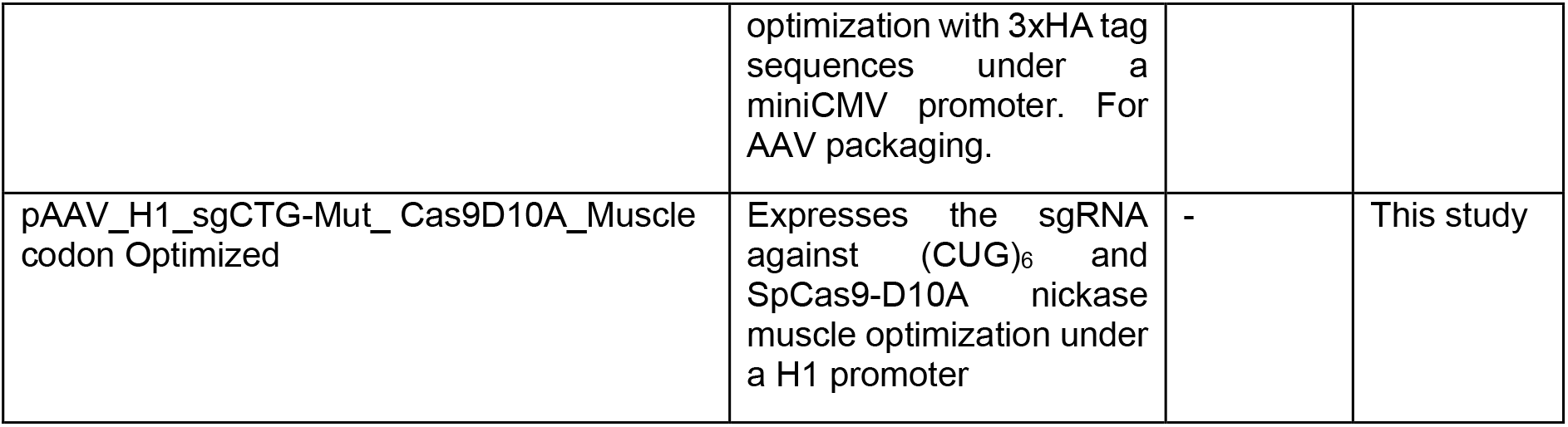
Plasmids used herein.

**Supplementary Table 4.**
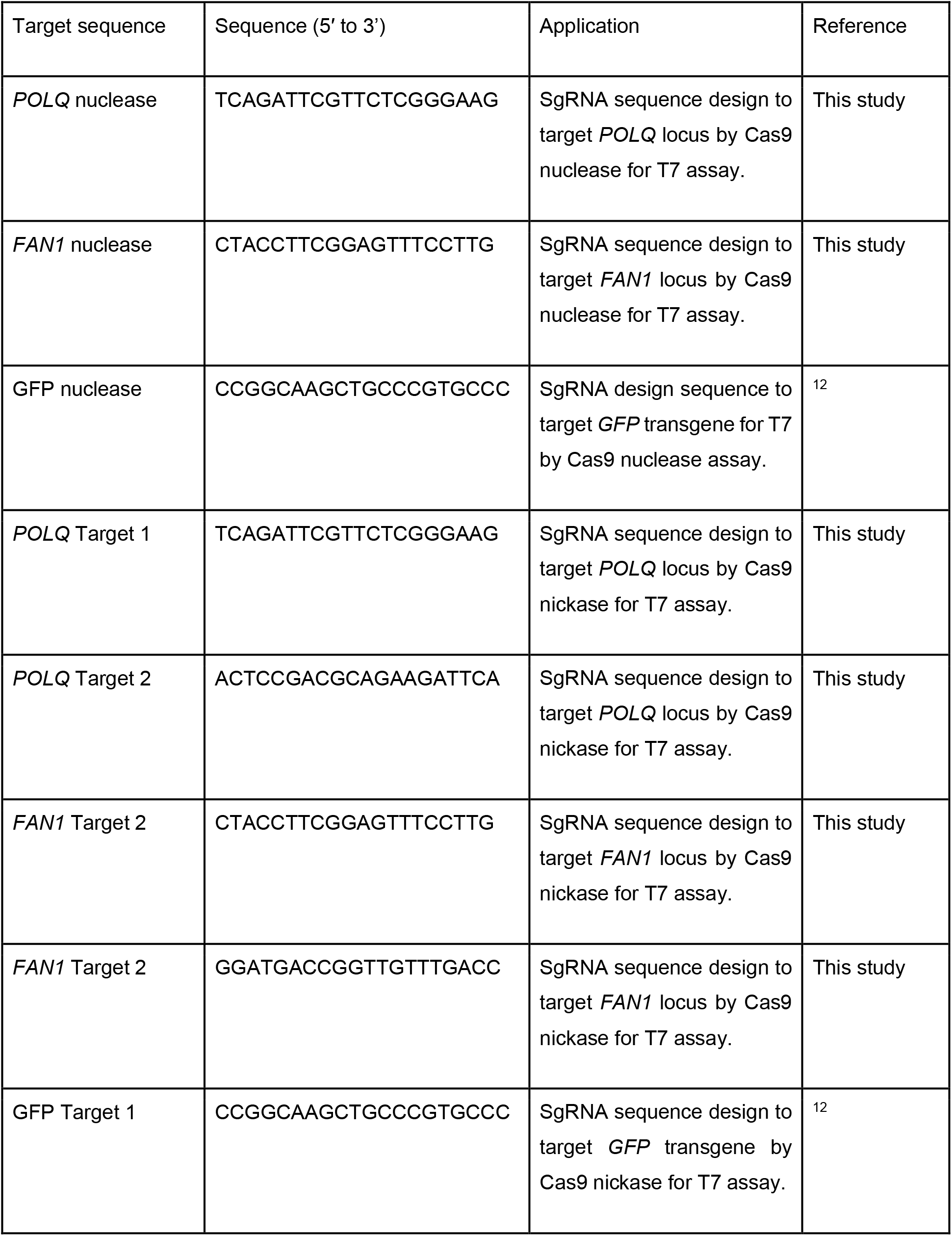
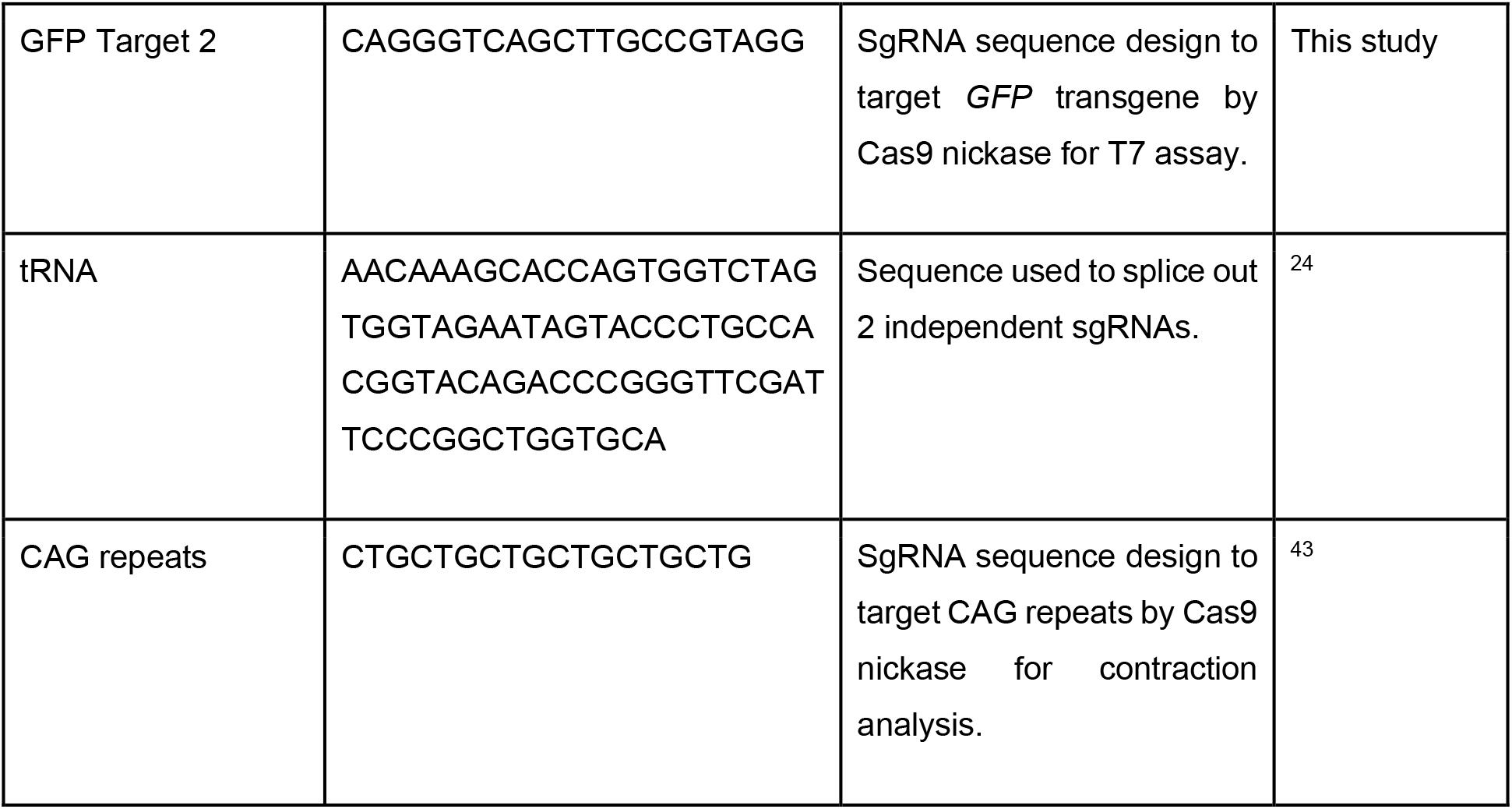
The sgRNA sequences used herein.

**Supplementary table 5:**
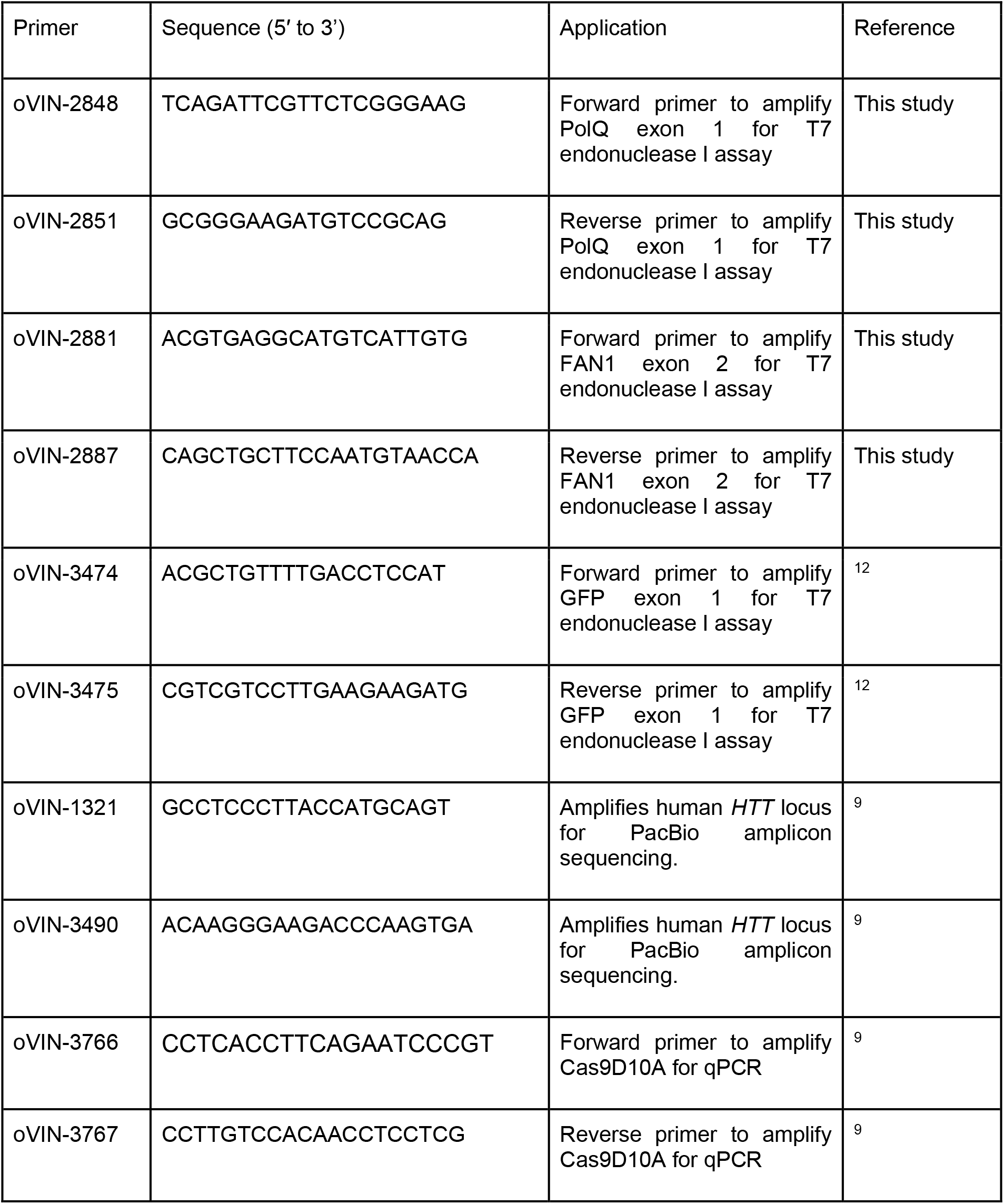
Primers used herein.

